# Inositol hexakisphosphate (IP6) accelerates immature HIV-1 Gag protein assembly towards kinetically-trapped morphologies

**DOI:** 10.1101/2022.03.29.486265

**Authors:** Alexander J. Pak, Manish Gupta, Mark Yeager, Gregory A. Voth

## Abstract

During the late stages of the HIV-1 lifecycle, immature virions are produced by the concerted activity of Gag polyproteins, primarily mediated by the capsid (CA) and spacer peptide 1 (SP1) domains, which assemble into a spherical lattice, package viral genomic RNA, and deform the plasma membrane. Recently, inositol hexakisphosphate (IP6) has been identified as an essential assembly cofactor that efficiently produces both immature virions *in vivo* and immature virus-like particles *in vitro*. To date, however, several distinct mechanistic roles for IP6 have been proposed on the basis of independent functional, structural, and kinetic studies. In this work, we investigate the molecular influence of IP6 on the structural outcomes and dynamics of CA/SP1 assembly using coarse-grained (CG) molecular dynamics (MD) simulations and free energy calculations. Here, we derive a bottom-up, low-resolution, and implicit-solvent CG model of CA/SP1 and IP6, and simulate their assembly under conditions that emulate both *in vitro* and *in vivo* systems. Our analysis identifies IP6 as an assembly accelerant that promotes curvature generation and fissure-like defects throughout the lattice. Our findings suggest that IP6 induces kinetically-trapped immature morphologies, which may be physiologically important for later stages of viral morphogenesis and potentially useful for virus-like particle technologies.

## Introduction

During the late stages of human immunodeficiency virus type 1 (HIV-1) replication, an immature particle is assembled at the plasma membrane interface and released^1-3^; this latter step is aided by the endosomal sorting complex required for transport (ESCRT) machinery.^1, 4^ The capsid (CA) and spacer peptide 1 (SP1) domains of the structural Gag polyprotein are responsible for coordinating Gag oligomerization into the immature lattice,^5-7^ which is both hexagonal and spherical (and therefore, incomplete).^7-10^ The matrix (MA) and nucleocapsid (NC) domains mediate interactions with the plasma membrane and viral genomic RNA, respectively.^1, 11-13^ The molecular mechanisms that regulate Gag assembly are potential targets for anti-retroviral therapies, as similarly demonstrated by capsid inhibitor drugs that perturb capsid assembly after maturation.^14-16^

The CA domain of Gag consists of two globular domains: the N-terminal (CA_NTD_) domain consists of 7 *α*-helices (helix 1-7) and the C-terminal (CA_CTD_) domain consists of 4 *α*-helices (helix 8-11).^17-19^ On the basis of atomistic structures resolved at 3.9 Å and 3.27 Å resolutions, using cryo-electron tomography (cryo-ET)^20^ and X-ray crystallography^21^, respectively, several critical inter-and intra-hexameric contacts have been identified. A trimeric contact formed by helix 2 stabilizes inter-hexameric interactions between CA_NTD_ while a dimeric contact formed by helix 1 stabilizes intra-hexameric interactions between CA_NTD_.^20, 22^ Inter-hexameric interactions are further mediated by the CA_CTD_ dimeric contact formed by helix 9, which is partially preserved in the mature CA configuration.^22-25^ Within the CA_CTD_ domain, the major homology region (MHR; residues I285-L304) loop and GVGG (residues G352-V355) β-turn are two structural motifs that contain important interactions, likely required to nucleate *α*-helical folding of the CA_CTD_/SP1 junction (residues P356-T371).^21^ This helical junction forms a six-helix bundle (6HB) that is stabilized by hydrophobic knobs-in-holes interactions, and represents an important intra-hexameric interaction that is minimally necessary for immature assembly.^21, 26^ Mutations within the aforementioned hexameric contacts have been shown to abrogate immature assembly.^6, 21, 26-27^

Beyond the protein-protein interactions described above, immature particle assembly requires other cellular constituents. For instance, viral RNA and to a lesser extent, generic oligonucleotides, serve as catalysts for protein multimerization by promoting colocalization^11, 28-30^; in the former case, this is primarily mediated by interactions between the NC domain and the RNA Ψ packaging signal.^31-32^ The plasma membrane serves as a scaffold to promote protein assembly through dimensional reduction,^11, 33-34^ which is directed by myristoyl insertion and interactions between the MA domain and phosphatidylinositol 4,5-bisphosphate (PIP_2_) lipids,^35-37^, and ultimately evolves into the lipid envelop of the released viral particle. *In vitro* assembly studies have previously identified short oligonucleotides and inositol hexakisphosphate (IP6) as two constituents minimally necessary for immature virus-like particle (VLPs) assembly.^30, 38-39^ In mammalian cells, IP6 is present in the cytoplasm at 10-40 µM concentrations.^40-41^ Recently, combined biochemical and structural studies have shown that IP6 is an essential cofactor that promotes immature particle production around a stoichiometric ratio of one IP6 molecule per immature hexamer.^42-43^ Within the immature hexamer, the negatively charged IP6 molecule binds between two six-membered lysine rings, K290 and K359, which line the interior pore of the hexamer and are positioned directly above the 6HB.^42^

While it is evident that the presence of IP6 is essential, the precise role of IP6 during immature virus assembly remains unclear. For example, is the role of IP6 to stabilize the 6HB by neutralizing the two lysine rings along the central pore? Previous *in vitro* studies have shown that CA/SP1 morphology is sensitive to solution pH or salt (e.g. NaCl or KCl) concentrations, which modulate electrostatic interactions^42, 44^; under high salt or low pH conditions, CA/SP1 proteins assemble into mature tubes while under low salt or high pH conditions, these same proteins assemble into spherical VLPs.^42, 44-45^ Intriguingly, alanine mutations of the lysine rings tended to increase *in vitro* VLP formation^46^ while reducing *in vivo* particle production,^43^ both with respect to wild-type (WT) virus. Alternatively, is the role of IP6 to influence assembly kinetics? For example, a recent study by Kucharska et al. suggests that Gag encodes all of the necessary interactions for assembly and that IP6 and viral RNA act as assembly rate modulators.^47^ Finally, it is possible that IP6 serves both roles in a synergistic fashion.

In this work, we use coarse-grained (CG) molecular dynamics (MD) simulations to investigate the molecular influence of IP6 on CA/SP1 assembly. To do so, we derive a bottom-up, low-resolution, and implicit-solvent CG model of CA/SP1 and IP6, and implement a MD trajectory-based enhanced sampling approach. By quantifying 3D and 2D assembly behavior under three different buffer conditions (150 mM monovalent salt, 150 mM salt + IP6, and 400 mM salt), as well as free energy calculations, we show that IP6 primarily acts as an assembly accelerant. Immature lattices formed through the influence of IP6 assemble at faster rates compared to without, thereby promoting curvature generation and fissure-like defects throughout the lattice. Our analysis suggests that IP6 induces kinetically-trapped immature morphologies, which furthermore, may be physiologically important for later stages of viral morphogenesis or leveraged for efficient production of VLPs.

## Results

### Coarse-grained modeling and simulation

We first derived “bottom-up” implicit-solvent CG models for CA/SP1 (35 sites) and for IP6 (1 site), which were parameterized from AA MD trajectories (depicted in **Fig. 1A**). The two reference AA MD simulations were of a CA/SP1 18-mer, i.e. a hexamer surrounded by six partial hexamers, and the same 18-mer complexed with seven IP6 molecules positioned between the K290 and K359 rings in each hexamer pore; the first trajectory was used to derive the CA/SP1 CG model while the second trajectory was used to derive CG interactions between CA/SP1 and IP6. The CG Hamiltonian consists of four additive terms (see **Fig. 1A**) that respectively represent the excluded volume, flexibility, electrostatics, and short-range attraction of each molecule. Electrostatics are represented by a screened Yukawa potential; monovalent salt concentrations are implicitly modeled by adjusting the Debye length. Complete details on the mapping and parameterization procedure are provided in *Methods*.

**Figure 1:**
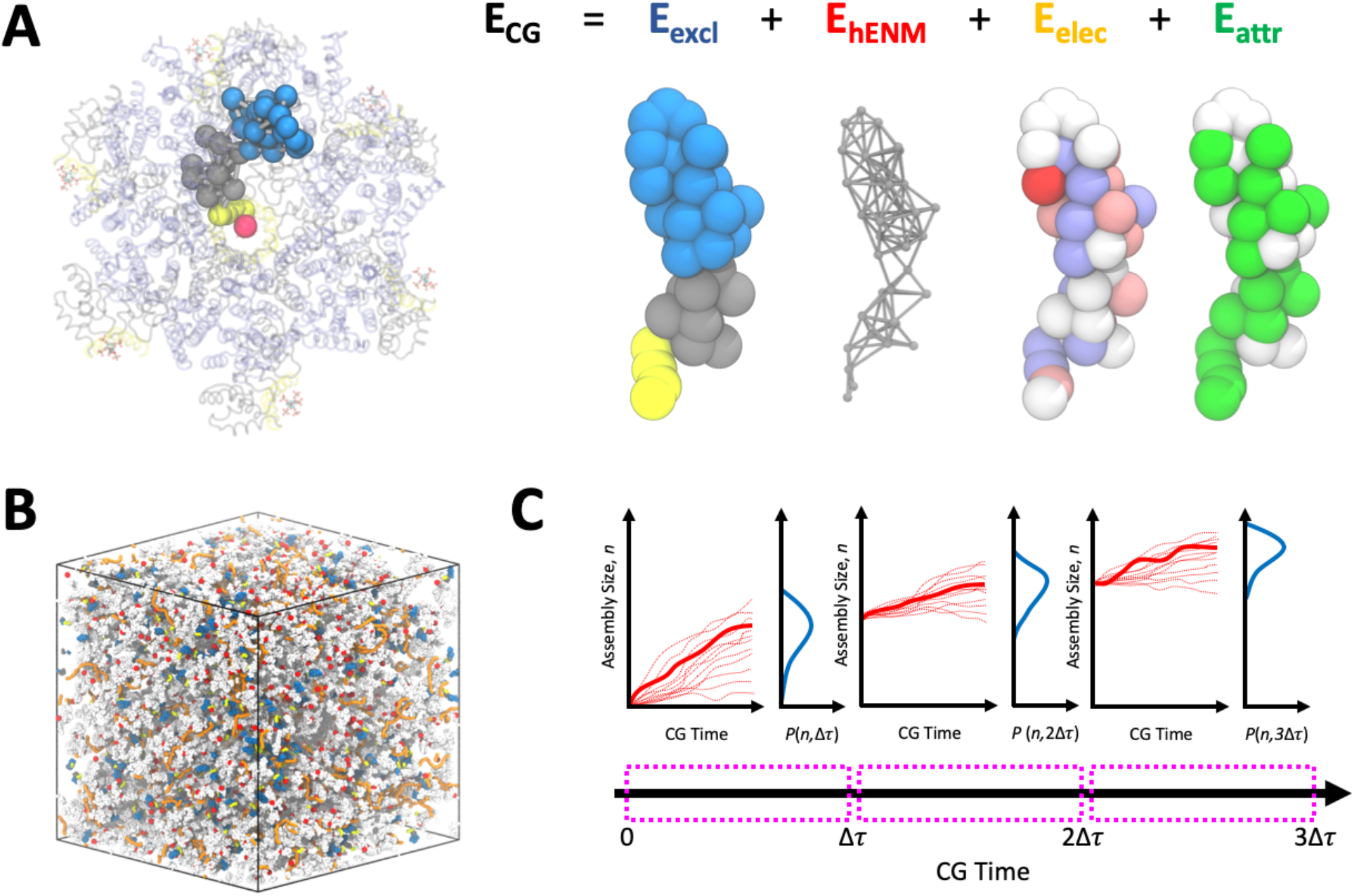
Schematic of coarse-grained (CG) HIV-1 CA/SP1 model and CGMD simulation. (A) Overlay of a CG CA/SP1 monomer (solid blue, grey, and yellow spheres) and IP6 monomer (red sphere) over an AA CA/SP1 18-mer (transparent blue, grey, and yellow tubes) bound to 7 IP6 molecules (cyan, purple, and red sticks); blue, grey, and yellow represent the NTD, CTD, and SP1 domains, respectively. The additive interaction elements of the CG Hamiltonian are depicted to the right; excluded volume (*E*_*excl*_), hetero-elastic network model (*E*_*hENM*_), screened Yukawa electrostatics (*E*_*elec*_), and protein-protein associative contacts (*E*_*attr*_). (B) A representative snapshot of an initial simulation configuration; active CA/SP1 (blue, grey, and yellow beads), inactive CA/SP1 (white beads), IP6 (red beads), and oligonucleotide (orange beads) molecules are randomly distributed. (C) Schematic of the parallel sampling technique used to propagate assembly during CGMD simulations. For each CGMD simulation of length Δ*τ*, multiple CGMD trajectories were spawned (thin red lines) from the previously selected trajectory. From the probability distribution of resultant assembly sizes (*P*(*n*)), a trajectory that corresponded to the maximum *P*(*n*) was selected to spawn the next generation of trajectories. All selected trajectories (thick red lines) were appended to create the final trajectory.

Using the aforementioned CG models, we performed two types of CG MD assembly simulations. We first simulated CA/SP1 assembly under conditions that emulate prior *in vitro* studies. Here, we included CA/SP1 with 50-nt oligonucleotide (represented as a linear polymer) randomly dispersed within a cubic box (see **Fig. 1B**) at (i) 150 mM monovalent salt, (ii) 400 mM monovalent salt, and (iii) 150 mM monovalent salt and IP6 conditions. We next simulated CA/SP1 under simplified *in vivo* conditions, i.e., at a plasma membrane interface. We used the same salt and IP6 conditions, and additionally included a 9000-nt oligonucleotide and a CG lipid membrane using a generic model with 30 *k*_*B*_*T* bending rigidity.

For both sets of simulations, we implemented a constant protein concentration algorithm and a path sampling algorithm. In the former, we intermittently identified CA/SP1 that have oligomerized with a hexameric CA/SP1 seed, which served as a nucleator. All remaining CA/SP1 were randomly partitioned into “active” and “inactive” monomers, similar to the ultra-coarse-graining strategy used in prior mature CA assembly simulations.^16, 48^ Assuming an excess of initial CA/SP1 is provided, this strategy ensures that a constant concentration of CA/SP1 is available for assembly. To aid assembly path sampling, we discretized the assembly process into sequential segments of length Δ*τ*_*CG*_ CG time steps. For each segment, 10 parallel CG MD simulations following randomized Langevin dynamics were propagated. Cluster sizes were computed at the final configuration of each parallel run, and the probability distribution of cluster sizes (*P*(*n*)) was approximated as a log-normal distribution. The trajectory yielding a cluster closest to the mean of *P*(*n*) was used to initialize the next set of parallel trajectories; the selected trajectory from each chuck was concatenated to create the final assembly trajectory, a strategy conceptually similar to parallel cascade MD.^49^ Since CA/SP1 assembly is a rare-event process, our strategy facilitates cluster formation and extracts trajectories that represent the mean kinetics predicted by the underlying CG model. Complete details on the CG MD simulations are provided in *Methods*.

To quantify free energies of monomer-monomer association, we performed 2D well-tempered metadynamics (WTMetaD) simulations.^50-51^ Each system contained two monomers that were biased against two collective variables (CVs): the distances between the two CA_CTD_ centroids and the two CA_NTD_ centroids; when included, IP6 was restrained to be proximal to the K290 and K359 residues of one monomer. To remove volumetric entropic effects, the free energies were shifted by the Jacobian correction factor *k*_*B*_*T* In *R*^2^.^52-53^ All reported free energy surfaces (FESs) were averaged over three independent replicas after alignment to the free energy at large distances. Additional details on the WTMetaD simulations are provided in *Methods*.

### Assembly into virus-like particles

*In vitro* experiments have shown that the presence of 500 µM IP6 induces CA/SP1 (also 500 µM) assembly into spherical VLPs at pH 6 and physiological salt conditions, while the absence of IP6 induces amorphous protein aggregates. Under high salt conditions, CA/SP1 instead assembles into tubules. Mutating K290 and K359 to alanine was previously shown to yield VLPs, suggesting that electrostatic repulsion of the lysine ring inhibits assembly, which may be recovered by salt-induced electrostatic screening. Here, we use CG MD simulations to investigate the differences between IP6-induced and salt-screening-induced CA/SP1 assembly under similar *in vitro* conditions.

We simulated 850 µM CA/SP1 assembly under three different conditions: (i) 150 mM monovalent salt (i.e., low salt), (ii) 400 mM monovalent salt (i.e., high salt), and (iii) 150 mM monovalent salt with 425 µM IP6; in all three cases, 50-nt RNA-representing polymers were also added to the system. Our results are summarized in **Fig. 2**. The time-series profiles in **Fig. 2A** show that while CA/SP1 fails to assemble at low-salt conditions, both high-salt and low-salt with IP6 conditions lead to productive protein assembly. We note that under our simulated salt and IP6 concentrations, the presence of IP6 (elevated salt) yields a lattice consisting of 999±89 (534±27) CA/SP1 after 1500×10^6^ *τ*_CG_, suggesting that IP6 accelerates CA/SP1 by nearly two-fold compared to high-salt conditions. Beyond accelerated kinetics, the presence of IP6 appears to shift the morphological outcome of CA/SP1 assembly. As depicted in **Fig. 2B**, CA/SP1 assembles into a spherical VLP under low-salt with IP6 conditions. Here, the particle consists of fissure defects that are characteristic of immature HIV-1 virions as seen in prior cryo-ET lattice maps.^3, 8, 54^ Alternatively, CA/SP1 assembles into a curved and contiguous hexameric lattice under high-salt conditions, as evident from **Fig. 2C**. Here, each protomer appears to maximize its inter-hexameric and intra-hexameric contacts, suggesting that the assembled lattice is representative of an energetically minimized state. In contrast, the VLP formed under the presence of IP6 is likely a kinetically trapped state that results from enhanced assembly kinetics. Prior assembly simulations have shown that similar defects emerge when protomer association energetics are enhanced ^11^, suggesting that IP6 interactions induce a comparable effect.

**Figure 2:**
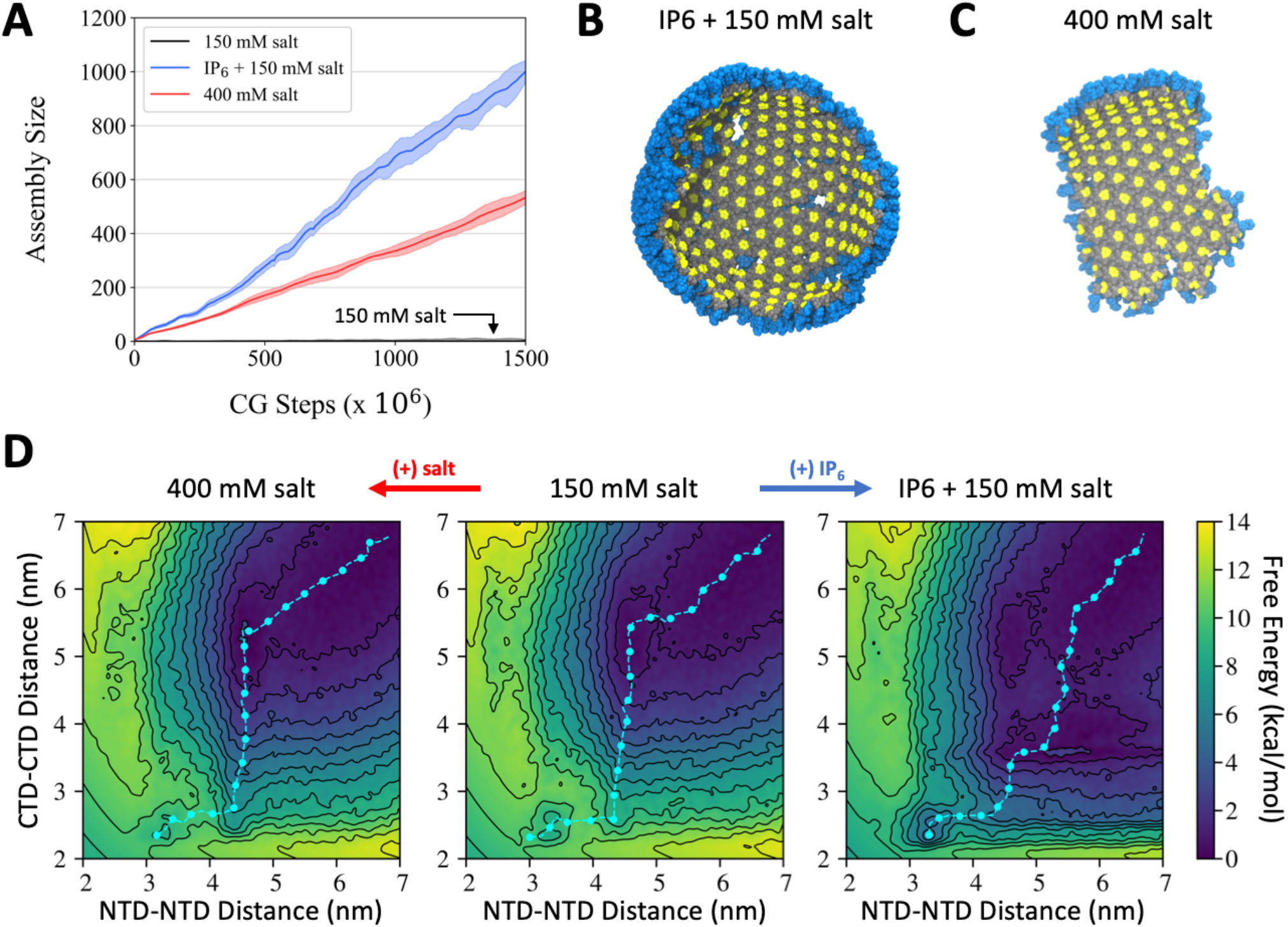
CGMD simulations of “*in vitro*” HIV-1 CA/SP1 assembly. (A) Time-series profiles of assembled protein cluster size as a function of CG time step under varying salt and co-factor conditions. (B) Snapshot of assembled protein under the presence of IP6 and 150 mM salt; blue, grey, and yellow balls represent the NTD, CTD, and SP1 domains of CA/SP1 while IP6 and RNA are not depicted for clarity. (C) Snapshot of assembled protein under the presence of 400 mM salt; same color scheme as (B). (D) Comparison of free energy profiles for the association of two CA/SP1 monomers projected along their inter-protein CTD and NTD distances under varying salt and co-factor conditions. The cyan points and line trace the minimum free energy path.

To quantify thermodynamic differences between protomer association under the aforementioned conditions, we performed WTMetaD simulations. We compare our computed FESs in **Fig. 2D**, which are projected onto the distances between CTD-CTD and NTD-NTD centers-of-mass. The low-salt FES serves as a baseline to characterize how the underlying FES is altered under high-salt and low-salt with IP6 conditions; recall that 850 µM CA/SP1 does not spontaneously assemble under these conditions (see Fig. 2A). All three cases exhibit qualitatively similar minimum free energy paths (MFEPs), in which association begins at the CTD-CTD interface, which is followed by additional association at both CTD-CTD and NTD-NTD interfaces. However, there are several key differences to note. For example, CTD-CTD association is facilitated under high-salt conditions, as demonstrated by the broadening of the FES towards smaller CTD-CTD distances when the NTD-NTD distance is below 4.5 nm. The shapes of the free energy minimum where the CTD-CTD (NTD-NTD) distance is around 2.5 (3.3) nm are also distinct. This local minimum is comparatively shallow in the absence of IP6, whereas the minimum is both deepened and broadened when IP6 is present. With IP6 present, the free energy barriers for CA/SP1 association and dissociation are 5.7 ± 0.6 and 1.5 ± 0.8 kcal/mol, respectively. The association barrier is notably lower than that of both low-salt (10.6 ± 0.1 kcal/mol) and high-salt (9.6 ± 0.2 kcal/mol) conditions, while the dissociation barriers (1.9 ± 0.3 and 0.8 ± 0.4 kcal/mol for low-salt and high-salt, respectively) are comparable across all three cases. In other words, rather than stabilizing the bound state, i.e., increasing the dissociation barrier, IP6 appears to promote CA/SP1 association by reducing the association barrier, in part by promoting co-localization up to CTD-CTD and NTD-NTD distances of 3.5 and 4.5 nm, respectively. We should note, however, that the above calculations only pertain to monomer-monomer association, while oligomer-monomer association may include additional collective effects. Nonetheless, these calculations serve as a minimal basis to identify and compare the impact of electrostatic screening and IP6 interactions on CA/SP1 assembly.

### Immature lattice assembly at the membrane

In our prior CG MD simulations of CA/SP1 assembly on a membrane, we had found that initial protein assembly first required a perturbation or “puncta” to the local membrane curvature^11^ as the assembling immature Gag without it could not sufficiently drive the membrane bending towards a budding event. Interestingly, prior experimental studies have shown that the number of virions released from cells is reduced by around 60-90% when IP6-producing enzymes are knocked out without loss of integrity to the produced virion.^43^ We therefore hypothesize that the effect of IP6 on the assembling Gag may induce the necessary membrane curvature generation (and therefore, force) necessary for productive immature lattice assembly and budding.

We thus simulated CA/SP1 assembly with RNA at a lipid membrane interface (30 *k*_*B*_*T* bending rigidity) under the same three conditions as above: (i) 150 mM monovalent salt (i.e. low salt), (ii) 400 mM monovalent salt (i.e. high salt), and (iii) 150 mM monovalent salt with IP6 (using a 1:1 CA/SP1:IP6 stoichiometric ratio); here, a 9000-nt RNA-representing polymer was added to the system. To reduce the lag time arising from the nucleation event, we ran short simulations until a small hexamer was formed, which we used to seed the rest of the lattice. We maintained a constant surface coverage of 2380 #/µm^2^ for CA/SP1 protein.

We summarize assembly time-series statistics in **Fig. 3A**. At our simulated protein concentrations, we find that CA/SP1 is unable to assemble at 150 mM salt; as discussed later, assembly under these salt concentrations is possible with elevated protein expression levels. In contrast, both 400 mM salt and 150 mM salt with IP6 conditions results in immature lattice growth. However, in the case of 150 mM salt with IP6, it is notable that the lattice grows to one containing 220±15 protomers after 400×10^6^ CG time steps, while the 400 mM salt case grows to one containing 90±20 protomers within the same simulation time. In other words, the presence of IP6 at 150 mM salt accelerates assembly nearly 2.5-fold compared to that of elevated salt at 400 mM concentrations (i.e. increased electrostatic screening). As seen in **Fig. 3B**, accelerated assembly kinetics under the presence of IP6 also appears to incorporate fissure-like defects throughout the lattice, which correspondingly results in greater curvature generation. In comparison, as seen in **Fig. 3C**, the lattice assembled under 400 mM salt remains uniform, contiguous, and flatter in comparison to the IP6-containing lattice. Although these assembly rates are slower than those computed for the *in vitro* system, we note that the two systems should not be directly compared due to differences in effective protein concentration and dimensional reduction.

**Figure 3:**
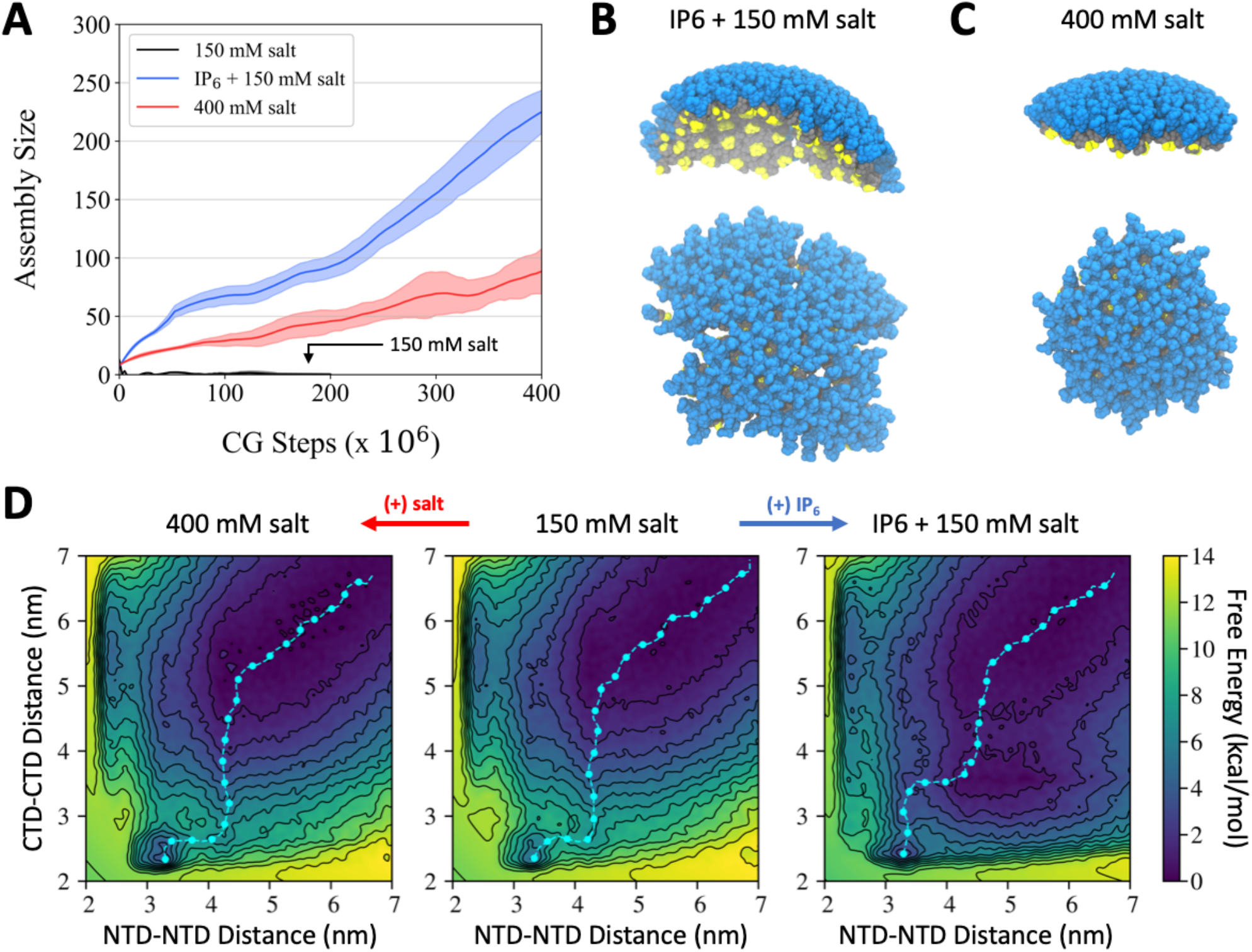
CGMD simulations of HIV-1 CA/SP1 assembly at a membrane interface. (A) Time-series profiles of assembled protein cluster size as a function of CG time step under varying salt and co-factor conditions. (B) Snapshot of assembled protein under the presence of IP6 and 150 mM salt; blue, grey, and yellow balls represent the NTD, CTD, and SP1 domains of CA/SP1 while IP6, RNA, and lipids are not depicted for clarity. (C) Snapshot of assembled protein under the presence of 400 mM salt; same color scheme as (B). (D) Comparison of free energy profiles for the association of two CA/SP1 monomers projected along their inter-protein CTD and NTD distances under varying salt and co-factor conditions. The cyan points and line trace the minimum free energy path.

In **Fig. 3D**, we quantify the 2D FES for CA/SP1 association when diffusion is restricted to the lateral *xy* plane. Similar to our prior analysis under *in vitro* conditions, we investigate the changes to the FES under high-salt and low-salt with IP6 conditions in comparison to low-salt conditions, under which extended assembly is not observed. Under high-salt conditions, the shape of the close contact free energy minimum located when the NTD-NTD and CTD-CTD distances are around 3.2 and 2.4 nm, respectively, is qualitatively similar to that of low-salt conditions. Interestingly, the free energy barriers for dissociation under low-salt and high-salt conditions are comparable (3.2 ± 0.6 and 4.1 ± 0.4 kcal/mol, respectively) while the barriers for association are slightly lowered in the latter case (7.9 ± 0.4 and 7.1 ± 0.3 kcal/mol, respectively, with *p* = 0.04 according to the unpaired t-test). Although the position of the free energy minimum is conserved under low-salt with IP6 conditions, the shape of this minimum is widened to favor an alternative CTD-CTD association path, while the barriers for dissociation and association are 2.0 ± 0.6 and 4.1 ± 0.4 kcal/mol, respectively. Hence, while the presence of IP6 yields a comparable dissociation barrier to that of the low-salt case, the association barrier is notably reduced by 3.8 ± 0.5 kcal/mol. It is also interesting to note that the association barriers across the three cases are consistently lower than that of the above *in vitro* system, while the dissociation barriers are consistently larger. Both effects are likely due to dimensional reduction, which effectively reduces the number of accessible transition state configurations.

To further quantify CA/SP1 assembly kinetics, we performed several shorter simulations (10×10^6^ CG MD timesteps) at varying protein surface coverage levels. While our previous simulations predict that CA/SP1 does not assemble under low-salt conditions at 2380 #/µm^2^ protein surface coverage, assembly may proceed at elevated concentrations. In particular, we depict the mean CA/SP1 assembly rate as a function of protein surface coverage and assembly conditions in **Fig. 4A**. At low surface coverage (≤ 2500 µm^-2^), both high-salt and low-salt with IP6 conditions promote assembly while low-salt conditions do not. At high surface coverage (≥ 2500 µm^-2^), all three conditions result in productive assembly, with the largest rates achieved under low-salt with IP6 conditions, followed by high-salt and low-salt conditions. Assuming power law rate kinetics, we find that all three assembly conditions follow sublinear scaling with respect to protein surface coverage, in which low-salt, high-salt, and low-salt with IP6 conditions have scaling law orders of 0.63, 0.66, and 0.74, respectively. At this point, it is not clear why sublinear reaction orders emerge. The impact of RNA as a substrate for CA/SP1 co-localization or the assembly of CA/SP1 into small intermediates prior to lattice growth,^55-58^ including the basic trimer-of-dimer unit that has been implicated for both mature and immature lattice assembly,^11, 48, 59^ are two factors that are likely to contribute. In the latter case, a sublinear reaction order may indicate that competing intermediates, that is, intermediates that are not productive for lattice assembly, increasingly form at elevated protein concentrations. Nonetheless, it is clear that the addition of IP6 and increased electrostatic screening due to increased salt consistently increase assembly rates compared to physiological salt conditions; low-salt with IP6 and high-salt conditions represent a 3-fold and 2-fold increase, respectively, in effective protein concentration compared to that of low-salt conditions.

**Figure 4.**
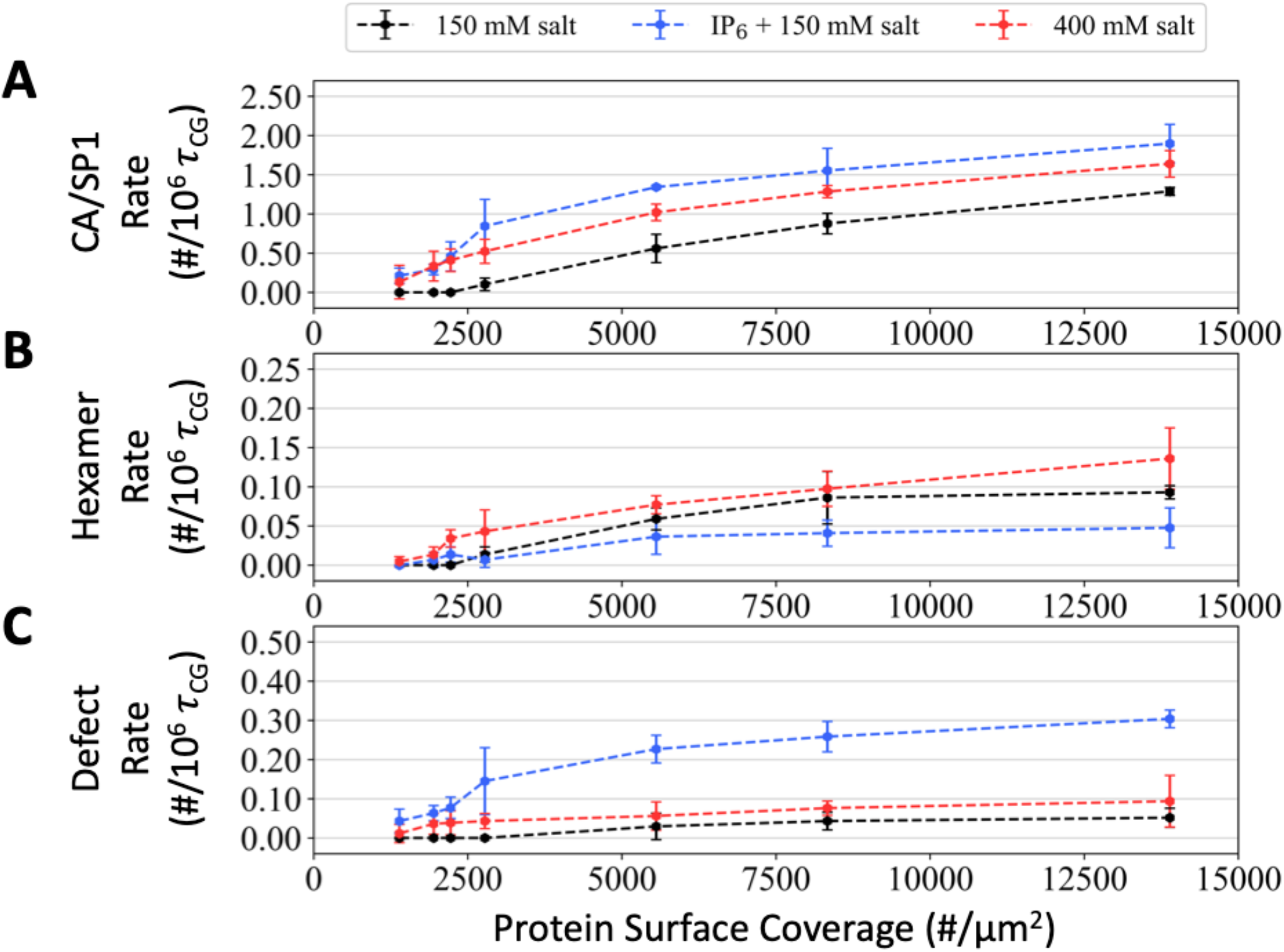
CA/SP1 assembly statistics under varying protein surface coverage densities. (A) Mean assembly rate (# CA/SP1 per 10^6^ CG timestep) describing the growth of the immature lattice under varying salt and co-factor conditions. (B,C) The mean formation rate (per 10^6^ CG timestep) for (B) hexamers and (C) defects (i.e. incomplete hexamer) within the immature lattice.

We further assess the morphological characteristics of the assembled lattices under varying protein surface coverages and assembly conditions. In **Fig. 4B** and **Fig. 4C**, we depict the rate of formation for complete and defective (i.e., incomplete) hexamers, respectively, within the assembled lattice. We find that high-salt conditions consistently yield the largest rate for hexamer growth while low-salt with IP6 conditions yield the lowest. Intriguingly, both high-salt and low-salt conditions accumulate defective hexamers at rates comparable to that of their hexamer growth rates, although reduced by around 30-50%. The inclusion of IP6, on the other hand, dramatically increases the rate of defect growth by 6-fold compared to the rate of hexamer growth. It is therefore evident that while defects are incorporated into the lattice under all three assembly conditions, and more so at elevated protein concentrations, the rate of defect incorporation is substantially increased due to the presence of IP6.

## Discussion

We have developed a bottom-up (i.e., derived from atomistic simulation data), low-resolution, and implicit-solvent CG model for HIV-1 CA/SP1 and IP6 in order to study their assembly behavior on length-and time-scales relevant to the formation of viral particles. Despite the simplicity of the model, our CGMD simulations recapitulate key experimental observations, including the selective inducement of spherical VLPs under the presence of IP6^42, 46^ that are morphologically consistent with immature lattices observed *in vivo*.^3, 8^ Furthermore, we have investigated the sensitivity of CA/SP1 assembly kinetics and structural outcomes to salt conditions and the presence of IP6.

Thermodynamically-driven self-assembly often favors a self-regulated nucleation and growth mechanism that ensures error correction and tends to consist of weak multivalent interactions between specific protomer-protomer interfaces resulting in the formation of structures that maximize these contacts.^60-61^ Assembly can also proceed along kinetically-trapped pathways that may yield morphologies alternative to the thermodynamically preferred structure. Our simulations suggest that Gag tubule formation, as seen under select *in vitro* conditions^44^ or with single point mutations,^62^ is slow and proceeds through disassembly/reassembly, a known error correction mechanism,^63^ whereas spherical particle formation is kinetically-driven, in which lattice defects represent kinetically-trapped states. In the latter case, the lattice grows too quickly for error correction by local remodeling, producing defects that subsequently facilitate curvature throughout the lattice. Thus, we propose that tubules are a near-equilibrium assembly product while VLPs are a kinetically-trapped product. Relatedly, we find that IP6 promotes kinetically-trapped morphologies under conditions where CA/SP1 would not otherwise assemble by lowering the free energy barrier for association rather than stabilizing the associated state, which accelerates CA/SP1 assembly.

The robust creation of immature lattice defects by IP6 may serve additional purposes beyond simple curvature generation. The ability to transform spherical particles into cone-shaped cores via Gag proteolysis is an intriguing property of HIV-1 maturation.^1^ During this process, viral protease (PR) cleaves Gag precursors at five different sites, in which cleavage of the site within the CA/SP1 6HB is the rate limiting step.^1, 64-65^ Our prior cryo-ET and MD study showed that lattice defects facilitate uncoiling of the CA/SP1 partial helical bundle, which is crucial for protease access to the CA/SP1 cleavage site.^54^ By inducing fissure-like defects via a kinetically-driven process, IP6 may facilitate the emergence of cleavage initiation sites during maturation.

Recent studies of Gag multimerization at the plasma membrane interface suggest that the extent of membrane deformation,^11^ and relatedly lipid phase segregation,^66^ are coupled to the dynamics of Gag assembly. On the basis of our membrane-bound simulations, we propose that the free energy penalty for lattice growth due to membrane deformation resistance is mitigated by the IP6-induced lattice defects, which serve to relieve strain from lattice curvature. In addition, our free energy calculations indicate that IP6 lowers the free energy of the protomer-protomer bound state by 2.6 ± 0.5 kcal/mol compared to the case when IP6 is absent, suggesting that IP6 lowers the free energy of formation for the lattice. Both factors may explain why IP6 appears to be essential for efficient virion production,^43^ although additional work is necessary to explain why other inositol phosphate molecules, such as inositol pentakisphosphate (IP5), and related polyanions are not as effective.

In the absence of IP6, CA/SP1 constructs have been shown to assemble into mature tubes *in vitro*^42, 46^ while our simulations produce relatively flatter and contiguous immature lattices. This observation underscores a limitation of the present CG model, which was trained using immature state statistics, and therefore restricted to the immature state. The mature state requires a structural transition from the immature state, which involves the uncoiling of the CA/SP1 helical junction and a hinge-like motion such that CA_NTD_ forms the primary hexameric pore.^24-25, 65, 67^ Incorporating this structural transition into future CG models is likely to be non-trivial, but may be aided by recent methodological efforts to recapitulate conformational transitions within CG space.^68^ Given that the presence of IP6 shifts mature CA assembly outcomes towards conical cores instead of tubules,^42^ we speculate that mature CA assembles into tubules following a near-equilibrium pathway while the presence of IP6 induces kinetically-driven assembly into mature cores. Nonetheless, the present work highlights the utility and importance of kinetically-driven assembly in the context of immature HIV-1 viral particle production, although additional studies are needed to understand the connection between the fissure-like defects, the kinetics of PR activity throughout the immature lattice, and mature CA assembly during the final stages of maturation.

In conclusion, our findings demonstrate the critical role of IP6 as both an assembly accelerant and morphological modulator of the immature lattice during the late stages of HIV-1 viral morphogenesis. Beyond HIV-1, IP6 has also been implicated as an assembly cofactor in other retroviruses, such as Rous sarcoma virus and equine infectious anemia virus.^69-70^ It would be interesting to investigate the utility of IP6 and its derivatives for the efficient production of broad VLPs that have been repurposed for biomedical technologies, including vaccine development and biologics delivery.^71-74^ Our results will hopefully stimulate future experimental studies as well as the use and development of even more sophisticated CG models.

## Methods

### All-Atom (AA) MD simulations

We used the 18-mer atomic model of CA/SP1 from Ref. 20 (PDB 5L93) as our initial protein structure. Seven IP6 molecules (each phosphate was deprotonated) were placed between the seven K290/K359 double lysine rings using Monte Carlo insertion. The protein-cofactor complex was solvated by 163010 water molecules, 646 Na^+^ ions, and 508 Cl^−^ ions (i.e. 150 mM NaCl) in a dodecahedron box with 1.5 nm of space between the edges of the protein and the box.

All simulations used the CHARMM36m force field^75^ and were performed using GROMACS 2016.^76^ Minimization was performed using steepest descent until the maximum force was reduced to 500 kJ/mol/nm. Then, equilibration was performed in several phases. First, 10 ns were integrated in the constant *NVT* ensemble at 310 K using the stochastic velocity rescaling thermostat^77^ with a damping time of 0.1 ps and a timestep of 2 fs. During this phase, the heavy atoms of the protein were harmonically restrained with a force constant of 1000 kJ/mol/nm^2^. An additional 10 ns were integrated in the constant *NPT* ensemble using the Nose-Hoover chain thermostat^78^ at 310 K (2 ps damping time) and the Parrinello-Rahman barostat^79^ at 1 bar (10 ps damping time) and a timestep of 2 fs; the restraints on the protein were maintained. Finally, an additional 530 ns were integrated in the constant *NVT* ensemble using the Nose-Hoover chain thermostat^78^ at 310 K (2 ps damping time) and a timestep of 2 fs. Throughout this procedure, H-containing bonds were constrained using the LINCS algorithm.^80^ Protein and cofactor configurations were gathered every 200 ps. The same process was repeated to generate a second trajectory without IP6 molecules. The final 500 ns of each trajectory were used for CG model generation.

### CG model generation

Using the generated AA MD trajectories described above, we derived a bottom-up model of CA/SP1 and IP6. To map the CA/SP1 protomer, we used the Essential Dynamics Coarse-Graining (EDCG) method^81^ to identify residue groupings that maximized overlap of the CG model motions with the AA collective motions. We specified 35 CG sites to balance computational efficiency and least-squared error in the represented principal component subspace. We mapped the IP6 molecule into a single CG site using a linear center-of-mass mapping.

After CG mapping, effective CG interactions (*E*_*CG*_) were determined using the following four terms:

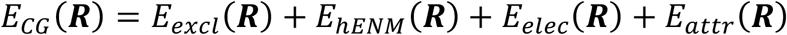

where *E*_*excl*_ represents excluded volume interactions, *E*_*hENM*_ represents intra-protein interactions, *E*_*elec*_ represents screened electrostatic interactions, *E*_*attr*_ represents attractive inter-protein interactions, and ***R*** represents configuration space of the CG coordinates. The hetero-elastic network model (hENM) method^82^ with bond energies *k*_*i*_ (*r*_*i*_ − *r* _*0,i*_)^2^, where *k*_*i*_ is the spring constant of a particular effective harmonic bond *i* and *r*_*0,i*_ is the equilibrium bond length for that bond, was used to represent *E*_*hENM*_. These parameters were optimized using the hENM method with a cutoff distance of 2 nm. For *E*_*excl*_, a soft cosine potential, 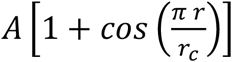, was used, where *A* = 25 kcal/mol and *r*_*c*_ is the onset for excluded volume determined from pair correlation functions. For *E*_*elec*_, a Yukawa potential,^83^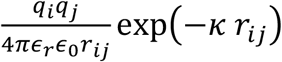, was used, where *q*_*i*_ is the aggregate charge of CG site *i, κ* = 1.274 or 0.481 nm^-1^ is the inverse Debye length for 150 and 400 mM NaCl, respectively, and *ε*_*r*_ is the effective dielectric constant of the protein environment, approximated as 17.5.^84^

Our CG model initially assumes that all close contacts between protomers contribute to protein association through two-body attractive interactions, i.e., *E*_*attr*_. We identified relevant CG site pairs on the basis of their pair correlation functions. If the first peak of the pair correlation function was located below 1.75 nm, and the standard deviation less than 0.15 nm, that pair of CG sites was selected. We represented *E*_*attr*_ as a pairwise Gaussian potential, 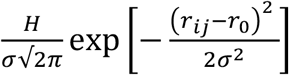, where *r*_*0*_ and *σ* are the mean and standard deviation determined by a fit to the first peak of the pair correlation function between CG sites *i* and *j* through least-squares regression. The constant *H* was optimized for each pair interaction using relative-entropy minimization (REM).^85^ During this process, some of the parameters exhibited diverging behavior or yielded repulsive interactions. Interactions with diverging behavior were removed from the CG model while the remaining parameters were reoptimized, although only minor changes were observed.

We used the same interaction parameters for the RNA polymer as in our prior CG study of HIV-1 Gag assembly^11^ and for the CG lipid as in our prior CG study of the SARS-CoV-2 virion.^86^

### CG assembly simulations

All simulations were prepared using PACKMOL and Moltemplate.^87-88^ For the *in vitro* simulations, 3012 CA/SP1 monomers, 200 50-nt RNA polymers, and 1500 IP6 molecules (when relevant) were randomly distributed throughout a 180×180×180 nm^3^ cubic box; two replicas were simulated. For the membrane simulations, a CG lipid bilayer was prepared using a grid distribution of 152352 lipids (76176 per monolayer) in a 450×450 nm^2^ lateral (*xy*) domain with an initial spacing of 4 nm between the two monolayer head groups in the *z* direction. An additional 4090 CA/SP1 monomers were uniformly distributed along a 2D grid 1.5 nm below the CG membrane followed by the 9000-nt RNA polymer (3 nm below the layer of CA/SP1) and a random distribution of 4096 IP6 molecules (3 nm below the RNA polymer). Due to computational cost, only one replica was simulated.

All CG MD simulations were performed using LAMMPS 18Jun2019.^89^ Conjugate gradient energy minimization was performed on each system until the change in force was less than 10^−6^. All systems were initially equilibrated using a Langevin thermostat^90^ at 310 K (5 ps damping constant) over 5×10^6^ *τ*_CG_ (with *τ*_CG_ = 50 fs) with attractive protein-protein interactions turned off to remove bias due to initial protein distributions. For the membrane simulations, a Berendsen barostat^91^ at 0 atm (10 ps damping constant, applied over the lateral *xy* dimension) was also applied; during this process, the *xy* simulation domain expanded to 454×454 nm^2^, which remained fixed for the remainder of the simulation.

We then integrated all CG molecules using a Langevin thermostat at 310 K (50 ps damping constant). We simultaneously integrated 10 trajectories over 50×10^6^ *τ*_CG_ with *τ*_CG_ = 50 fs; during these simulations, we maintained an 85% and 15% active population of CA/SP1 which were randomly selected every 0.5×10^6^ *τ*_CG_ for the *in vitro* and membrane-bound simulations, respectively. Trajectory snapshots were saved every 5×10^6^ *τ*_CG_. At the end of each set of 10 CGMD simulations, the sizes of the largest assembled CA/SP1 cluster in each trajectory was computed and their distribution fit to a log-normal function. The variance of the distribution was used to compute the standard error of the mean cluster size. The trajectory closest to the mean cluster size was used to initiate the next set of 10 simultaneous trajectories.

### CG metadynamics simulations

Free energy profiles were computed using the well-tempered metadynamics method^51^ via LAMMPS interfaced with the PLUMED 2.4 plugin.^92^ We used the same CGMD conditions as above. To emulate membrane-bound effects, we used two planar indenter forces (*F*_*indent*_(*z*) = *K*(*z* − *z*_0_)^2^) to restrict the *z*-diffusion of the NTD domain; here, *K* = 1 kcal/mol/Å^2^. Gaussian biases with a height of 0.1 kcal/mol and width of 0.05 nm were deposited every 1000 *τ*_CG_ using a bias factor of 2 *k*_*B*_*T*. The biases were deposited along two collective variables (CVs): (i) the distance between the centers-of-mass of the CTD and (ii) that of the NTD. A harmonic restraint was applied with a force constant of 1000 kcal/mol/Å to prevent the two CVs from exceeding 8 nm.

### CG analysis and visualization

Assembled clusters were identified using two proximity criteria. Inter-hexameric contacts were defined by the CTD dimer interface, i.e. when the distance between CG types 25/26 were less than 1.8 nm. Intra-hexameric contacts were defined by the top of the CA/SP1 6HB, i.e. when the distance between CG type 32 was less than 1.8 nm. For each trajectory snapshot, an adjacency graph was constructed and extracted using the NetworkX 2.1 (http://networkx.github.io/) Python package. Extracted clusters were visualized using VMD 1.9.3.^93^

## Acknowledgements

This research was supported by the National Institute of Allergy and Infectious Diseases (NIAID) of the National Institutes of Health under grant R01 AI154092. A.J.P gratefully acknowledges support from the National Institute of Allergy and Infectious Diseases of the National Institutes of Health under grant F32 AI150477. The authors also acknowledge the Texas Advanced Computing Center (TACC) at The University of Texas at Austin for providing HPC and visualization resources that have contributed to the research results reported within this paper.^94^ This work also used the Extreme Science and Engineering Discovery Environment (XSEDE), which is supported by the National Science Foundation grant number ACI-1548562.^95^

## Data Availability

Model parameters, simulation input files, and custom code are available on Github at: https://github.com/uchicago-voth/MSCG-models/tree/master/HIV_CASP1. The input files and data generated from this study are accessible from Zenodo: https://zenodo.org/record/6335601.

## Table of Contents (TOC) Graphic

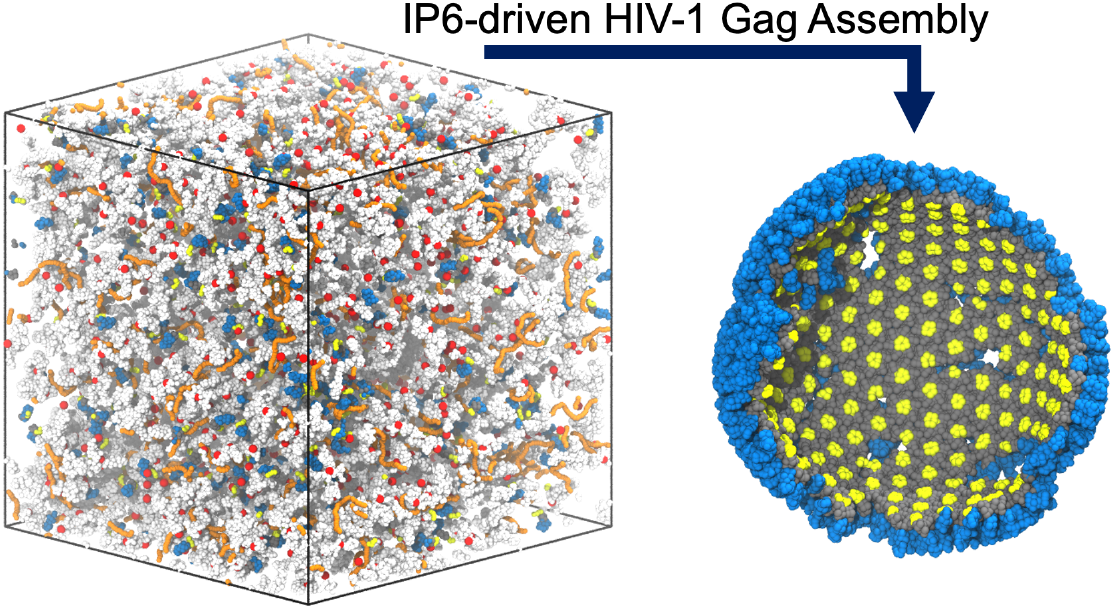

## References

1. Sundquist, W. I.; Kräusslich, H. G., HIV-1 assembly, budding, and maturation. Cold Spring Harb Perspect Med 2012, 2 (7), a006924.

2. Pornillos, O.; Ganser-Pornillos, B. K., Maturation of retroviruses. Curr Opin Virol 2019, 36, 47–55.

3. Briggs, J. A.; Riches, J. D.; Glass, B.; Bartonova, V.; Zanetti, G.; Krausslich, H. G., Structure and assembly of immature HIV. Proc Natl Acad Sci U S A 2009, 106 (27), 11090–5.

4. Popova, E.; Popov, S.; Gottlinger, H. G., Human immunodeficiency virus type 1 nucleocapsid p1 confers ESCRT pathway dependence. J Virol 2010, 84 (13), 6590–7.

5. Datta, S. A. K.; Clark, P. K.; Fan, L.; Ma, B.; Harvin, D. P.; Sowder, R. C.; Nussinov, R.; Wang, Y.-X.; Rein, A.; Sundquist, W. I., Dimerization of the SP1 Region of HIV-1 Gag Induces a Helical Conformation and Association into Helical Bundles: Implications for Particle Assembly. Journal of Virology 2016, 90 (4), 1773–1787.

6. Datta, S. A.; Temeselew, L. G.; Crist, R. M.; Soheilian, F.; Kamata, A.; Mirro, J.; Harvin, D.; Nagashima, K.; Cachau, R. E.; Rein, A., On the role of the SP1 domain in HIV-1 particle assembly: a molecular switch? J Virol 2011, 85 (9), 4111–21.

7. Wright, E. R.; Schooler, J. B.; Ding, H. J.; Kieffer, C.; Fillmore, C.; Sundquist, W. I.; Jensen, G. J., Electron cryotomography of immature HIV-1 virions reveals the structure of the CA and SP1 Gag shells. EMBO J 2007, 26 (8), 2218–26.

8. Carlson, L. A.; de Marco, A.; Oberwinkler, H.; Habermann, A.; Briggs, J. A.; Krausslich, H. G.; Grunewald, K., Cryo electron tomography of native HIV-1 budding sites. PLoS Pathog 2010, 6 (11), e1001173.

9. Johnson, D. S.; Bleck, M.; Simon, S. M., Timing of ESCRT-III protein recruitment and membrane scission during HIV-1 assembly. Elife 2018, 7.

10. Carlson, L. A.; Briggs, J. A.; Glass, B.; Riches, J. D.; Simon, M. N.; Johnson, M. C.; Muller, B.; Grunewald, K.; Krausslich, H. G., Three-dimensional analysis of budding sites and released virus suggests a revised model for HIV-1 morphogenesis. Cell Host Microbe 2008, 4 (6), 592–9.

11. Pak, A. J.; Grime, J. M. A.; Sengupta, P.; Chen, A. K.; Durumeric, A. E. P.; Srivastava, A.; Yeager, M.; Briggs, J. A. G.; Lippincott-Schwartz, J.; Voth, G. A., Immature HIV-1 lattice assembly dynamics are regulated by scaffolding from nucleic acid and the plasma membrane. Proc Natl Acad Sci 2017, 114 (47), E10056–E10065.

12. Muriaux, D.; Darlix, J. L., Properties and functions of the nucleocapsid protein in virus assembly. RNA Biol 2010, 7 (6), 744–53.

13. Yang, Y.; Qu, N.; Tan, J.; Rushdi, M. N.; Krueger, C. J.; Chen, A. K., Roles of Gag-RNA interactions in HIV-1 virus assembly deciphered by single-molecule localization microscopy. Proc Natl Acad Sci U S A 2018, 115 (26), 6721–6726.

14. Shi, J.; Zhou, J.; Shah, V. B.; Aiken, C.; Whitby, K., Small-molecule inhibition of human immunodeficiency virus type 1 infection by virus capsid destabilization. J Virol 2011, 85 (1), 542–9.

15. Bester, S. M.; Wei, G.; Zhao, H.; Adu-Ampratwum, D.; Iqbal, N.; Courouble, V. V.; Francis, C.; Annamalai, A. S.; Singh, P. K.; Shkriabai, N.; Van Blerkom, P.; Morrison, J.; Poeschla, E. M.; Engelman, A. N.; Melikyan, G. B.; Griffin, P. R.; Fuchs, J. R.; Asturias, F. J.; Kvaratskhelia, M., Structural and mechanistic bases for a potent HIV-1 capsid inhibitor. Science 2020, 370 (6514), 360–364.

16. Pak, A. J.; Grime, J. M. A.; Yu, A.; Voth, G. A., Off-Pathway Assembly: A Broad-Spectrum Mechanism of Action for Drugs That Undermine Controlled HIV-1 Viral Capsid Formation. J Am Chem Soc 2019, 141 (26), 10214–10224.

17. Du, S.; Betts, L.; Yang, R.; Shi, H.; Concel, J.; Ahn, J.; Aiken, C.; Zhang, P.; Yeh, J. I., Structure of the HIV-1 full-length capsid protein in a conformationally trapped unassembled state induced by small-molecule binding. J Mol Biol 2011, 406 (3), 371–86.

18. Momany, C.; Kovari, L. C.; Prongay, A. J.; Keller, W.; Gitti, R. K.; Lee, B. M.; Gorbalenya, A. E.; Tong, L.; McClure, J.; Ehrlich, L. S.; Summers, M. F.; Carter, C.; Rossmann, M. G., Crystal structure of dimeric HIV-1 capsid protein. Nat Struct Biol 1996, 3 (9), 763–70.

19. Gamble, T. R.; Yoo, S.; Vajdos, F. F.; von Schwedler, U. K.; Worthylake, D. K.; Wang, H.; McCutcheon, J. P.; Sundquist, W. I.; Hill, C. P., Structure of the carboxyl-terminal dimerization domain of the HIV-1 capsid protein. Science 1997, 278 (5339), 849–53.

20. Schur, F. K. M.; Obr, M.; Hagen, W. J. H.; Wan, W.; Jakobi, A. J.; Kirkpatrick, J. M.; Sachse, C.; Kräusslich, H.-G.; Briggs, J. A. G., An atomic model of HIV-1 capsid-SP1 reveals structures regulating assembly and maturation. Science 2016, 353, 506–508.

21. Wagner, J. M.; Zadrozny, K. K.; Chrustowicz, J.; Purdy, M. D.; Yeager, M.; Ganser-Pornillos, B. K.; Pornillos, O., Crystal structure of an HIV assembly and maturation switch. Elife 2016, 5.

22. Schur, F. K.; Hagen, W. J.; Rumlová, M.; Ruml, T.; Müller, B.; Kräusslich, H. G.; Briggs, J. A., Structure of the immature HIV-1 capsid in intact virus particles at 8.8 Å resolution. Nature 2015, 517 (7535), 505–8.

23. Zhao, G.; Perilla, J. R.; Yufenyuy, E. L.; Meng, X.; Chen, B.; Ning, J.; Ahn, J.; Gronenborn, A. M.; Schulten, K.; Aiken, C.; Zhang, P., Mature HIV-1 capsid structure by cryo-electron microscopy and all-atom molecular dynamics. Nature 2013, 497 (7451), 643–6.

24. Gres, A. T.; Kirby, K. A.; KewalRamani, V. N.; Tanner, J. J.; Pornillos, O.; Sarafianos, S. G., STRUCTURAL VIROLOGY. X-ray crystal structures of native HIV-1 capsid protein reveal conformational variability. Science 2015, 349 (6243), 99–103.

25. Pornillos, O.; Ganser-Pornillos, B. K.; Kelly, B. N.; Hua, Y.; Whitby, F. G.; Stout, C. D.; Sundquist, W. I.; Hill, C. P.; Yeager, M., X-ray structures of the hexameric building block of the HIV capsid. Cell 2009, 137 (7), 1282–92.

26. Accola, M. A.; Strack, B.; Göttlinger, H. G., Efficient particle production by minimal Gag constructs which retain the carboxy-terminal domain of human immunodeficiency virus type 1 capsid-p2 and a late assembly domain. J Virol 2000, 74 (12), 5395–402.

27. Guo, X.; Roldan, A.; Hu, J.; Wainberg, M. A.; Liang, C., Mutation of the SP1 sequence impairs both multimerization and membrane-binding activities of human immunodeficiency virus type 1 Gag. J Virol 2005, 79 (3), 1803–12.

28. Gross, I.; Hohenberg, H.; Krausslich, H. G., In vitro assembly properties of purified bacterially expressed capsid proteins of human immunodeficiency virus. Eur J Biochem 1997, 249 (2), 592–600.

29. Khorchid, A.; Halwani, R.; Wainberg, M. A.; Kleiman, L., Role of RNA in facilitating Gag/Gag-Pol interaction. J Virol 2002, 76 (8), 4131–7.

30. Muriaux, D.; Mirro, J.; Harvin, D.; Rein, A., RNA is a structural element in retrovirus particles. Proc Natl Acad Sci U S A 2001, 98 (9), 5246–51.

31. Webb, J. A.; Jones, C. P.; Parent, L. J.; Rouzina, I.; Musier-Forsyth, K., Distinct binding interactions of HIV-1 Gag to Psi and non-Psi RNAs: implications for viral genomic RNA packaging. RNA 2013, 19 (8), 1078–88.

32. Ding, P.; Kharytonchyk, S.; Waller, A.; Mbaekwe, U.; Basappa, S.; Kuo, N.; Frank, H. M.; Quasney, C.; Kidane, A.; Swanson, C.; Van, V.; Sarkar, M.; Cannistraci, E.; Chaudhary, R.; Flores, H.; Telesnitsky, A.; Summers, M. F., Identification of the initial nucleocapsid recognition element in the HIV-1 RNA packaging signal. Proc Natl Acad Sci U S A 2020, 117 (30), 17737–17746.

33. Dick, R. A.; Vogt, V. M., Membrane interaction of retroviral Gag proteins. Front Microbiol 2014, 5, 187.

34. Yandrapalli, N.; Muriaux, D.; Favard, C., Lipid domains in HIV-1 assembly. Front Microbiol 2014, 5, 220.

35. Monje-Galvan, V.; Voth, G. A., Binding mechanism of the matrix domain of HIV-1 gag on lipid membranes. Elife 2020, 9.

36. Vlach, J.; Saad, J. S., Structural and molecular determinants of HIV-1 Gag binding to the plasma membrane. Front Microbiol 2015, 6, 232.

37. Ono, A.; Ablan, S. D.; Lockett, S. J.; Nagashima, K.; Freed, E. O., Phosphatidylinositol (4,5) bisphosphate regulates HIV-1 gag targeting to the plasma membrane. P Natl Acad Sci USA 2004, 101 (41), 14889–14894.

38. Campbell, S.; Fisher, R. J.; Towler, E. M.; Fox, S.; Issaq, H. J.; Wolfe, T.; Phillips, L. R.; Rein, A., Modulation of HIV-like particle assembly in vitro by inositol phosphates. Proc Natl Acad Sci U S A 2001, 98 (19), 10875–9.

39. Campbell, S.; Vogt, V. M., Self-Assembly in-Vitro of Purified Ca-Nc Proteins from Rous-Sarcoma Virus and Human-Immunodeficiency-Virus Type-1. Journal of Virology 1995, 69 (10), 6487–6497.

40. Letcher, A. J.; Schell, M. J.; Irvine, R. F., Do mammals make all their own inositol hexakisphosphate? Biochem J 2008, 416 (2), 263–70.

41. Barker, C. J.; Wright, J.; Hughes, P. J.; Kirk, C. J.; Michell, R. H., Complex changes in cellular inositol phosphate complement accompany transit through the cell cycle. Biochemical Journal 2004, 380, 465–473.

42. Dick, R. A.; Zadrozny, K. K.; Xu, C.; Schur, F. K. M.; Lyddon, T. D.; Ricana, C. L.; Wagner, J. M.; Perilla, J. R.; Ganser-Pornillos, B. K.; Johnson, M. C.; Pornillos, O.; Vogt, V. M., Inositol phosphates are assembly co-factors for HIV-1. Nature 2018, 560 (7719), 509–512.

43. Mallery, D. L.; Faysal, K. M. R.; Kleinpeter, A.; Wilson, M. S. C.; Vaysburd, M.; Fletcher, A. J.; Novikova, M.; Böcking, T.; Freed, E. O.; Saiardi, A.; James, L. C., Cellular IP6 Levels Limit HIV Production while Viruses that Cannot Efficiently Package IP6 Are Attenuated for Infection and Replication. Cell Reports 2019, 29 (12), 3983-3996.e4.

44. Gross, I.; Hohenberg, H.; Wilk, T.; Wiegers, K.; Grattinger, M.; Muller, B.; Fuller, S.; Krausslich, H. G., A conformational switch controlling HIV-1 morphogenesis. EMBO J 2000, 19 (1), 103–13.

45. von Schwedler, U. K.; Stemmler, T. L.; Klishko, V. Y.; Li, S.; Albertine, K. H.; Davis, D. R.; Sundquist, W. I., Proteolytic refolding of the HIV-1 capsid protein amino-terminus facilitates viral core assembly. EMBO J 1998, 17 (6), 1555–68.

46. Dostalkova, A.; Kaufman, F.; Krizova, I.; Vokata, B.; Ruml, T.; Rumlova, M., In Vitro Quantification of the Effects of IP6 and Other Small Polyanions on Immature HIV-1 Particle Assembly and Core Stability. J Virol 2020, 94 (20).

47. Kucharska, I.; Ding, P.; Zadrozny, K. K.; Dick, R. A.; Summers, M. F.; Ganser-Pornillos, B. K.; Pornillos, O., Biochemical Reconstitution of HIV-1 Assembly and Maturation. J Virol 2020, 94 (5).

48. Grime, J. M. A.; Dama, J. F.; Ganser-Pornillos, B. K.; Woodward, C. L.; Jensen, G. J.; Yeager, M.; Voth, G. A., Coarse-grained simulation reveals key features of HIV-1 capsid self-assembly. Nat Commun 2016, 7, 11568.

49. Harada, R.; Kitao, A., Parallel Cascade Selection Molecular Dynamics (PaCS-MD) to generate conformational transition pathway. J Chem Phys 2013, 139 (3), 035103.

50. Dama, J. F.; Parrinello, M.; Voth, G. A., Well-tempered metadynamics converges asymptotically. Phys Rev Lett 2014, 112 (24), 240602.

51. Barducci, A.; Bussi, G.; Parrinello, M., Well-tempered metadynamics: a smoothly converging and tunable free-energy method. Phys Rev Lett 2008, 100 (2), 020603.

52. Boresch, S.; Karplus, M., The Jacobian factor in free energy simulations. Journal of Chemical Physics 1996, 105 (12), 5145–5154.

53. Sprik, M.; Ciccotti, G., Free energy from constrained molecular dynamics. Journal of Chemical Physics 1998, 109 (18), 7737–7744.

54. Tan, A.; Pak, A. J.; Morado, D. R.; Voth, G. A.; Briggs, J. A. G., Immature HIV-1 assembles from Gag dimers leaving partial hexamers at lattice edges as potential substrates for proteolytic maturation. Proceedings of the National Academy of Sciences 2021, 118 (3), e2020054118.

55. Morozov, A. Y.; Bruinsma, R. F.; Rudnick, J., Assembly of viruses and the pseudo-law of mass action. Journal of Chemical Physics 2009, 131 (15).

56. Zlotnick, A., Distinguishing reversible from irreversible virus capsid assembly. Journal of Molecular Biology 2007, 366 (1), 14–18.

57. Bruinsma, R. F.; Gelbart, W. M.; Reguera, D.; Rudnick, J.; Zandi, R., Viral self-assembly as a thermodynamic process. Physical Review Letters 2003, 90 (24).

58. Hagan, M. F.; Chandler, D., Dynamic pathways for viral capsid assembly. Biophysical Journal 2006, 91 (1), 42–54.

59. Tsiang, M.; Niedziela-Majka, A.; Hung, M.; Jin, D. B.; Hu, E.; Yant, S.; Samuel, D.; Liu, X. H.; Sakowicz, R., A Trimer of Dimers Is the Basic Building Block for Human Immunodeficiency Virus-1 Capsid Assembly. Biochemistry-Us 2012, 51 (22), 4416–4428.

60. Williams, R. J.; Smith, A. M.; Collins, R.; Hodson, N.; Das, A. K.; Ulijn, R. V., Enzyme-assisted self-assembly under thermodynamic control. Nat Nanotechnol 2009, 4 (1), 19–24.

61. Whitelam, S.; Jack, R. L., The statistical mechanics of dynamic pathways to self-assembly. Annu Rev Phys Chem 2015, 66, 143–63.

62. Bharat, T. A. M.; Menendez, L. R. C.; Hagen, W. J. H.; Lux, V.; Igonet, S.; Schorb, M.; Schur, F. K. M.; Krausslich, H. G.; Briggs, J. A. G., Cryo-electron microscopy of tubular arrays of HIV-1 Gag resolves structures essential for immature virus assembly. P Natl Acad Sci USA 2014, 111 (22), 8233–8238.

63. Keller, P. W.; Huang, R. K.; England, M. R.; Waki, K.; Cheng, N.; Heymann, J. B.; Craven, R. C.; Freed, E. O.; Steven, A. C., A two-pronged structural analysis of retroviral maturation indicates that core formation proceeds by a disassembly-reassembly pathway rather than a displacive transition. J Virol 2013, 87 (24), 13655–64.

64. Pettit, S. C.; Lindquist, J. N.; Kaplan, A. H.; Swanstrom, R., Processing sites in the human immunodeficiency virus type 1 (HIV-1) Gag-Pro-Pol precursor are cleaved by the viral protease at different rates. Retrovirology 2005, 2, 66.

65. Mattei, S.; Tan, A.; Glass, B.; Muller, B.; Krausslich, H. G.; Briggs, J. A. G., High-resolution structures of HIV-1 Gag cleavage mutants determine structural switch for virus maturation. P Natl Acad Sci USA 2018, 115 (40), E9401–E9410.

66. Sengupta, P.; Seo, A. Y.; Pasolli, H. A.; Song, Y. E.; Johnson, M.; Lippincott-Schwartz, J., A lipid-based partitioning mechanism for selective incorporation of proteins into membranes of HIV particles. Nat Cell Biol 2019, 21 (4), 452-+.

67. Wang, M. Z.; Quinn, C. M.; Perilla, J. R.; Zhang, H. L.; Shirra, R.; Hou, G. J.; Byeon, I. J.; Suiter, C. L.; Ablan, S.; Urano, E.; Nitz, T. J.; Aiken, C.; Freed, E. O.; Zhang, P. J.; Schulten, K.; Gronenborn, A. M.; Polenova, T., Quenching protein dynamics interferes with HIV capsid maturation. Nature Communications 2017, 8.

68. Sharp, M. E.; Vazquez, F. X.; Wagner, J. W.; Dannenhoffer-Lafage, T.; Voth, G. A., Multiconfigurational Coarse-Grained Molecular Dynamics. Journal of Chemical Theory and Computation 2019, 15 (5), 3306–3315.

69. Dick, R. A.; Xu, C. Y.; Morado, D. R.; Kravchuk, V.; Ricana, C. L.; Lyddon, T. D.; Broad, A. M.; Feathers, J. R.; Johnson, M. C.; Vogt, V. M.; Perilla, J. R.; Briggs, J. A. G.; Schur, F. K. M., Structures of immature EIAV Gag lattices reveal a conserved role for IP6 in lentivirus assembly. Plos Pathogens 2020, 16 (1).

70. Obr, M.; Ricana, C. L.; Nikulin, N.; Feathers, J. P. R.; Klanschnig, M.; Thader, A.; Johnson, M. C.; Vogt, V. M.; Schur, F. K. M.; Dick, R. A., Structure of the mature Rous sarcoma virus lattice reveals a role for IP6 in the formation of the capsid hexamer. Nature Communications 2021, 12 (1).

71. Ding, X. W.; Liu, D.; Booth, G.; Gao, W.; Lu, Y., Virus-Like Particle Engineering: From Rational Design to Versatile Applications. Biotechnol J 2018, 13 (5).

72. Frietze, K. M.; Peabody, D. S.; Chackerian, B., Engineering virus-like particles as vaccine platforms. Current Opinion in Virology 2016, 18, 44–49.

73. Hill, B. D.; Zak, A.; Khera, E.; Wen, F., Engineering Virus-like Particles for Antigen and Drug Delivery. Curr Protein Pept Sc 2018, 19 (1), 112–127.

74. Banskota, S.; Raguram, A.; Suh, S.; Du, S. W.; Davis, J. R.; Choi, E. H.; Wang, X.; Nielsen, S. C.; Newby, G. A.; Randolph, P. B.; Osborn, M. J.; Musunuru, K.; Palczewski, K.; Liu, D. R., Engineered virus-like particles for efficient in vivo delivery of therapeutic proteins. Cell 2022, 185 (2), 250–265.

75. Huang, J.; Rauscher, S.; Nawrocki, G.; Ran, T.; Feig, M.; de Groot, B. L.; Grubmüller, H.; MacKerell Jr, A. D., CHARMM36m: an improved force field for folded and intrinsically disordered proteins. Nature Methods 2016, 14, 71.

76. Abraham, M. J.; Murtola, T.; Schulz, R.; Páll, S.; Smith, J. C.; Hess, B.; Lindahl, E., GROMACS: High performance molecular simulations through multi-level parallelism from laptops to supercomputers. SoftwareX 2015, 1-2, 19–25.

77. Bussi, G.; Zykova-Timan, T.; Parrinello, M., Isothermal-isobaric molecular dynamics using stochastic velocity rescaling. The Journal of Chemical Physics 2009, 130 (7), 074101.

78. Martyna, G. J.; Klein, M. L.; Tuckerman, M., Nosé–Hoover chains: The canonical ensemble via continuous dynamics. The Journal of Chemical Physics 1992, 97 (4), 2635–2643.

79. Parrinello, M.; Rahman, A., Crystal Structure and Pair Potentials: A Molecular-Dynamics Study. Phys Rev Lett 1980, 45 (14), 1196–1199.

80. Hess, B.; Bekker, H.; Berendsen, H. J. C.; Fraaije, J. G. E. M., LINCS: A linear constraint solver for molecular simulations. Journal of Computational Chemistry 1997, 18 (12), 1463–1472.

81. Zhang, Z.; Lu, L.; Noid, W. G.; Krishna, V.; Pfaendtner, J.; Voth, G. A., A Systematic Methodology for Defining Coarse-Grained Sites in Large Biomolecules. Biophysical Journal 2008, 95 (11), 5073–5083.

82. Lyman, E.; Pfaendtner, J.; Voth, G. A., Systematic Multiscale Parameterization of Heterogeneous Elastic Network Models of Proteins. Biophysical Journal 2008, 95 (9), 4183–4192.

83. Yukawa, H., On the Interaction of Elementary Particles. I. Proceedings of the Physico-Mathematical Society of Japan. 3rd Series 1935, 17, 48–57.

84. Li, L.; Li, C.; Zhang, Z.; Alexov, E., On the Dielectric “Constant” of Proteins: Smooth Dielectric Function for Macromolecular Modeling and Its Implementation in DelPhi. Journal of Chemical Theory and Computation 2013, 9 (4), 2126–2136.

85. Shell, M. S., The relative entropy is fundamental to multiscale and inverse thermodynamic problems. The Journal of Chemical Physics 2008, 129 (14), 144108.

86. Yu, A.; Pak, A. J.; He, P.; Monje-Galvan, V.; Casalino, L.; Gaieb, Z.; Dommer, A. C.; Amaro, R. E.; Voth, G. A., A multiscale coarse-grained model of the SARS-CoV-2 virion. Biophys J 2021, 120 (6), 1097–1104.

87. Martínez, L.; Andrade, R.; Birgin, E. G.; Martínez, J. M., PACKMOL: A package for building initial configurations for molecular dynamics simulations. Journal of Computational Chemistry 2009, 30 (13), 2157–2164.

88. Jewett, A. I.; Zhuang, Z.; Shea, J.-E., Moltemplate a Coarse-Grained Model Assembly Tool. Biophysical Journal 2013, 104 (2), 169a.

89. Plimpton, S., Fast Parallel Algorithms for Short-Range Molecular Dynamics. Journal of Computational Physics 1995, 117 (1), 1–19.

90. Vanden-Eijnden, E.; Ciccotti, G., Second-order integrators for Langevin equations with holonomic constraints. Chemical Physics Letters 2006, 429 (1), 310–316.

91. Berendsen, H. J. C.; Postma, J. P. M.; van Gunsteren, W. F.; DiNola, A.; Haak, J. R., Molecular dynamics with coupling to an external bath. The Journal of Chemical Physics 1984, 81 (8), 3684–3690.

92. Tribello, G. A.; Bonomi, M.; Branduardi, D.; Camilloni, C.; Bussi, G., PLUMED 2: New feathers for an old bird. Comput Phys Commun 2014, 185 (2), 604–613.

93. Humphrey, W.; Dalke, A.; Schulten, K., VMD: visual molecular dynamics. J Mol Graph 1996, 14 (1), 33-8, 27-8.

94. Stanzione, D.; West, J.; Evans, R. T.; Minyard, T.; Ghattas, O.; Panda, D. K. In Frontera: The Evolution of Leadership Computing at the National Science Foundation, Practice and Experience in Advanced Research Computing (PEARC ‘20), July 26, 2020; 2020; pp 106–111.

95. Towns, J.; Cockerill, T.; Dahan, M.; Foster, I.; Gaither, K.; Grimshaw, A.; Hazlewood, V.; Lathrop, S.; Lifka, D.; Peterson, G. D.; Roskies, R.; Scott, J. R.; Wilkins-Diehr, N., XSEDE: Accelerating Scientific Discovery. Computing in Science & Engineering 2014, 16 (5), 62–74.

